# Modular Combinatorial Binding among Human *Trans*-acting Factors Reveals Direct and Indirect Factor Binding

**DOI:** 10.1101/027953

**Authors:** Yuchun Guo, David K. Gifford

## Abstract

The combinatorial binding of *trans*-acting factors (TFs) to regulatory genomic regions is an important basis for the spatial and temporal specificity of gene regulation. We present a new computational approach that reveals how TFs are organized into combinatorial regulatory programs. We define a regulatory program to be a set of TFs that bind together at a regulatory region. Unlike other approaches to characterizing TF binding, we permit a regulatory region to be bound by one or more regulatory programs. We have developed a method called regulatory program discovery (RPD) that produces compact and coherent regulatory programs from *in vivo* binding data using a topic model. Using RPD we find that the binding of 115 TFs in K562 cells can be organized into 49 interpretable regulatory programs that bind ~140,000 distinct regulatory regions in a modular manner. The discovered regulatory programs recapitulate many published protein-protein physical interactions and have consistent functional annotations of chromatin states. We found that, for certain TFs, direct (motif present) and indirect (motif absent) binding is characterized by distinct sets of binding partners and that the binding of other TFs can predict whether the TF binds directly or indirectly with high accuracy. Joint analysis across two cell types reveals both cell-type-specific and shared regulatory programs and that thousands of regulatory regions use different programs in different cell types. Overall, our results provide comprehensive cell-type-specific combinatorial binding maps and suggest a modular organization of binding programs in regulatory regions.

## Introduction

The combinatorial binding of *trans*-acting factors (TFs) is an important basis for the spatial and temporal specificity of gene regulation (Georges et al. 2010; Spitz and Furlong 2012; Weingarten-Gabbay and Segal 2014). Combinations of TFs have been shown to regulate gene expression stripes in the Drosophila embryo (Stanojevic et al. 1991), to generate cell-type-specific signaling responses (Mullen et al. 2011; Trompouki et al. 2011), and to program cell fates (Mazzoni et al. 2013). However, previous studies have examined only a limited number of well-known factors. Here we perform a systematic and unbiased dissection of the combinatorial binding patterns of 115 TFs to reveal their interactions and roles in gene regulation in selected human cell types. Our analysis method discovers sets of TFs that bind together to the same regulatory regions, and we call each distinct set of co-binding TFs a *regulatory program*. The systematic discovery of regulatory programs has recently become possible with large-scale efforts such as the ENCODE project to comprehensively profile the *in vivo* binding of tens to hundreds of TFs in multiple cell types (The ENCODE Project Consortium 2012).

Gene regulation is initiated by the interaction of enhancer-bound TFs, promoter-bound TFs, and TFs that bring the enhancers and promoters together in three dimensions. Previous studies have found that TFs tend to bind in clusters, which are typically characterized by a large number of TF binding sites in a regulatory region (Gerstein et al. 2012; Yip et al. 2012; Gerstein et al. 2010; modENCODE Consortium et al. 2010). We will represent complex factor binding patterns as the co-occurrence of multiple binding programs with diverse functions in the same genomic regions. For example, since CTCF and cohesin can be physically associated with both enhancers and promoters, a CTCF/cohesin program may co-occur with enhancer-related programs, promoter-related programs, or both (Guo et al. 2012b; Phillips-Cremins et al. 2013). Hence, understanding the interplay among regulatory programs is important to dissect the complexity of gene regulation.

Previous methods have modeled TF binding at a given regulatory region with only a single regulatory program. For example, self-organizing maps (SOMs) have been used to explore and visualize the colocalization patterns of TFs (Xie et al. 2013); k-means clustering has been used to characterize combinatorial regulation of erythroid enhancers (Xu et al. 2012). These hard-clustering-based methods assign each regulatory region to at most one program, and are thus not suitable for modeling co-binding by multiple regulatory programs with diverse functions. Non-negative matrix factorization (NMF), a soft clustering method, has also been applied to infer TF interactions (Giannopoulou and Elemento 2013). However, this work did not explicitly explore the issue of multiple program usage and predicted only a small number of TF combinations. It notably failed to capture the well-studied CTCF/cohesin interaction (Phillips-Cremins et al. 2013; Rubio et al. 2008).

A probabilistic topic model is a mixed-membership model that can represent modular regulatory program usage in regulatory regions. Topic models have been widely used to discover thematic structures in a large corpus of documents (Blei 2012; Blei et al. 2003). A topic model decomposes documents into a set of all shared topics, where a topic is a set of words that co-occur in multiple documents. Each document is described by a set of topics that occur in the document, which allows document exploration and organization. The factoring of a document into multiple topics permits the discovery of compact and coherent topics that can be combined to accurately represent a document. This factoring results in better performance in predicting held-out data than mixture models,(Blei et al. 2003)(Blei et al. 2003)(Blei et al. 2003) as a mixture model forces a document to be described by a single topic (Blei et al. 2003). Topic modeling has been used to discover gene expression programs (Bicego et al. 2010; Gerber et al. 2007) and microRNA regulatory modules (Joung and Fei 2009), but it has not yet been applied to study TF combinatorial binding.

We present a new computational method called regulatory program discovery (RPD) that applies a topic model to systematically discover the regulatory programs using a large compendium of *in vivo* TF binding data. Applying RPD to data from human K562 and GM12878 cells, we discovered diverse sets of regulatory programs and found that tens of thousands of regulatory regions use multiple programs in a modular manner. We found that, for certain TFs, direct (motif present) and indirect (motif absent) binding is characterized by distinct sets of binding partners. Joint analysis across two cell types reveals cell-type-specific and shared programs, and that thousands of regulatory regions use different programs in different cell types. Overall, our results provide comprehensive cell-type-specific global combinatorial binding maps and suggest a modular organization of binding programs in regulatory regions.

## Results

### Regulatory program discovery (RPD) discovers compact and coherent regulatory programs

RPD discovers regulatory programs given binding data for a large set of regulatory regions. RPD is based on Hierarchical Dirichlet Processes (Teh et al. 2006), a Bayesian non-parametric topic model that automatically determines the number of programs based on the complexity of the observed data. To use conventional document topic model terminology, regulatory regions are “documents,” TF binding sites are “words,” and regulatory programs are “topics.” As in the document model, a regulatory region may utilize one or more regulatory programs, and a TF may participate in multiple regulatory programs.

The co-binding of TFs in regulatory regions across the genome can be represented without loss by a region-TF matrix. In the work below, a full region-TF matrix would be of size ~140,000 regions by 115 TFs and is difficult to directly interpret. Using topic modeling, a large region-TF matrix is summarized into a compact program-TF matrix (49 programs x 115 TFs) and a region-program assignment table. The program-TF matrix describes the set of regulatory programs discovered, with each program represented as a probability distribution over all the TFs. The assignment table assigns each TF binding site from each region to one of the programs. The assignment table can be further summarized into a region-program matrix (~140,000 regions x 49 programs) that describes which regulatory programs each regulatory region uses (Fig. 1A).

**Figure 1.**
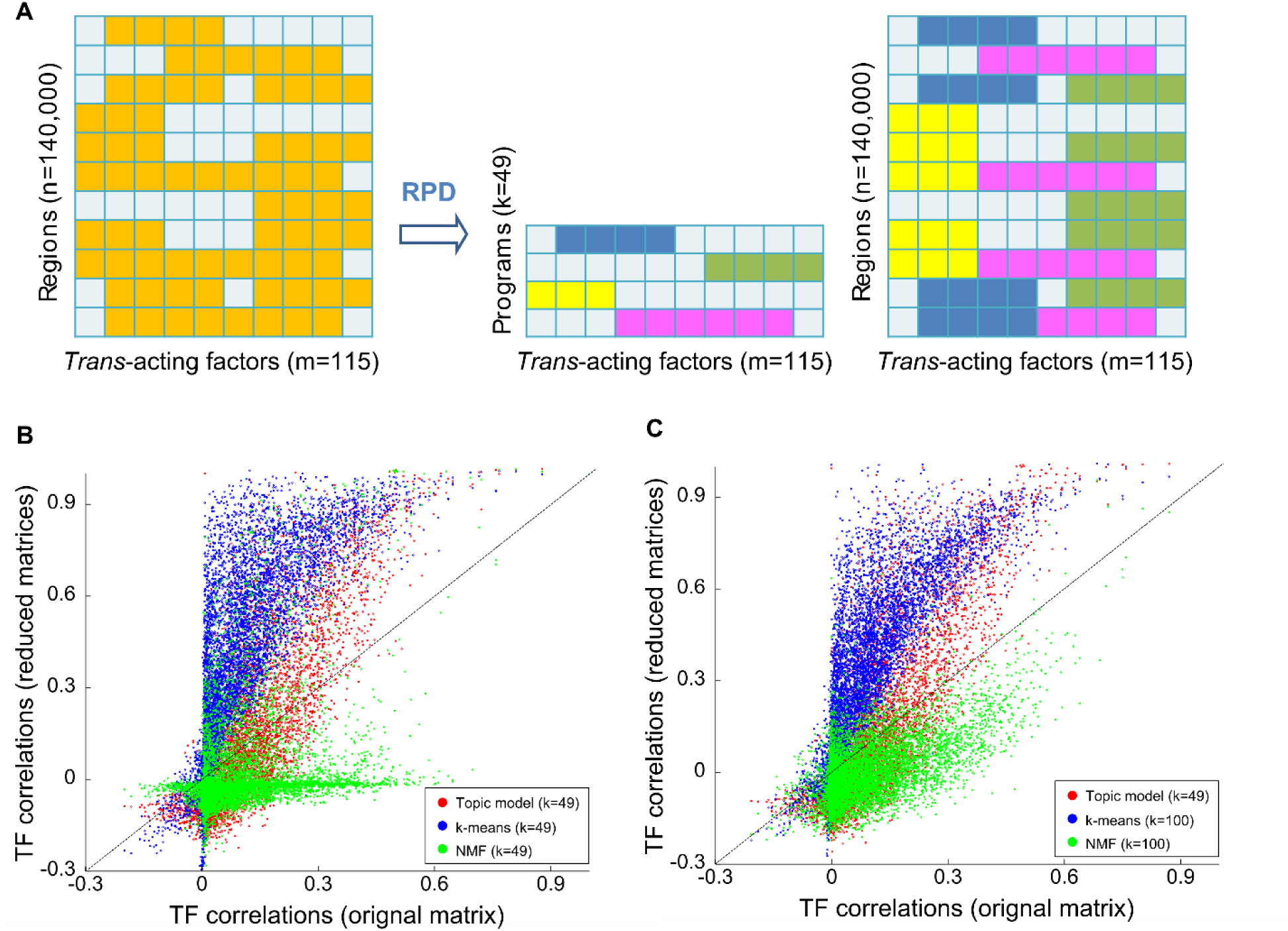
Regulatory program discovery (RPD) discovers compact and coherent regulatory programs. A) RPD discovers *trans*-acting factor (TF) combinatorial binding patterns (regulatory programs, shown in various colors) from a large number of discrete regions across the genome. A region may use one or more programs. A factor may participate in one or more programs. RPD uses a topic model to summarize a large compendium of binding data into a small number of regulatory programs and to assign binding sites to the programs. B) The topic model re-capitulates the original binding data more accurately than k-means clustering and non-negative matrix factorization (NMF). Each point in the scatter plot represents the Pearson correlation coefficient between a pair of TFs calculated using the original binding data (x-axis) or calculated using the reduced data matrix (k=49) by topic model, k-means clustering, or NMF (y-axis). C) The scatter plot is similar to B), but with k=100 for k-means clustering and NMF.

We first tested the ability of RPD, k-means clustering, and NMF to accurately capture TF-TF correlations that are present in their input binding data. We applied all three approaches to the same set of K562 cell ChIP-seq binding data to discover the same number of programs (k=49). We then evaluated how well the pairwise TF correlation scores from the program-TF matrix derived from these three methods correlate to the pairwise TF correlation scores from the original region-TF matrix. We found that the programs learned from the topic model more accurately recapitulate the pairwise TF co-binding relationship in the original data (r=0.81) than the programs learned from the other two methods (r=0.74 for k-means clustering, r=0.30 for NMF) (Fig. 1B). When k is increased to 100, the performance of k-means clustering and NMF improves to r=0.80 and r=0.59, respectively (Fig. 1C). Increasing k to the total number of regions will exactly recapitulate the original binding data but will not permit common patterns of binding to be observed. Similarly, applying a SOM to the same dataset from K562 cells partitioned the regulatory regions into several thousand of co-binding patterns (Xie et al. 2013). The increase in program count makes the interpretation of the programs harder.

We then asked if the RPD programs were a better representation of modular factor binding than the clusters produced by k-means. For this analysis, we considered an RPD program to be matched to a k-means cluster and vice-versa if the Pearson correlation for their TF site count distributions is greater than 0.5. We found that all the k-means clusters (k=49 and k=100) are matched by RPD programs, while 10 RPD programs are not matched by any k-means cluster for k=49, and 6 RPD programs are not matched by any k-means cluster when k is 100 (Supplemental Fig. S1). The RPD programs not matched to k-means clusters are a consequence of RPD’s ability to represent binding patterns that not discovered by clustering. Clustering may not discover these programs because they are used by relatively fewer regulatory regions and often co-occur with other programs.

In summary, our analysis shows that RPD is better at capturing the modular combinatorial binding of TFs and learning a set of compact and coherent programs than hard clustering (or similar approaches such as self-organizing maps) and matrix factorization approaches.

### RPD discovers a compact set of human regulatory programs

We next discovered regulatory programs in a single cell type using a compendium of ChIP-seq data from 115 TFs in human K562 cells (The ENCODE Project Consortium 2012). All the TF binding data were pooled and merged into ~140,000 non-overlapping co-binding regions, each of which was constrained to contain at least 3 TF binding sites. Applying RPD, we discovered a global combinatorial binding map consisting of 49 regulatory programs (RPs) (Fig. 2, Supplemental Table S1). We also applied PRD to data (86 TFs) from GM12878 cells and discovered 49 programs (Supplemental Fig. S2, Supplemental Table S1).

**Figure 2.**
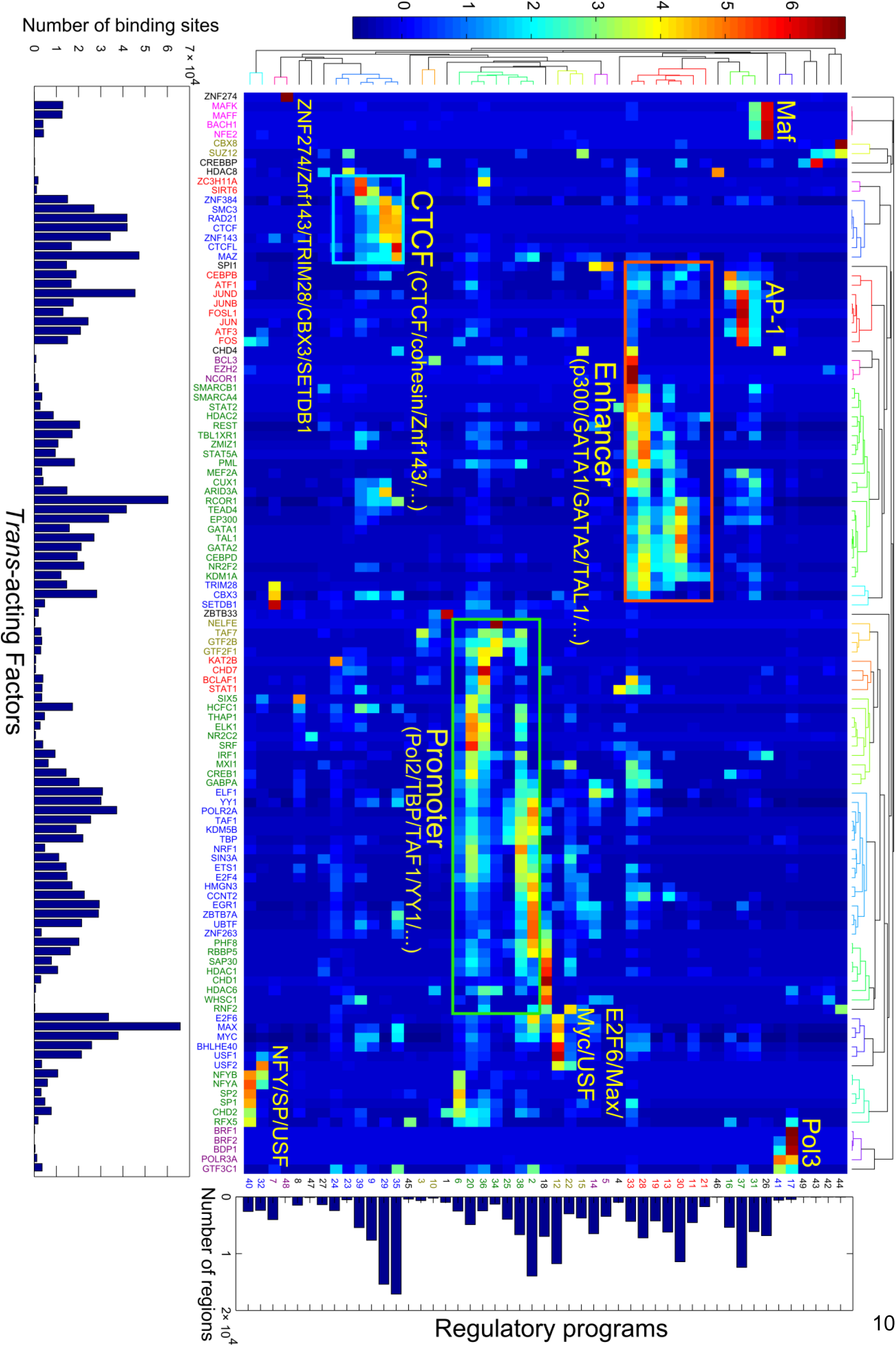
*Trans*-acting factors bind in a complex combinatorial and context-specific manner. RPD was applied to a compendium of ChIP-seq binding sites of 115 TFs (~140,000 regions) in human K562 cells and discovered 49 regulatory programs. Each cell in the heatmap represents the z-score of the TF binding site count (standardized along the columns) of a TF (column) in a regulatory program (row). The bottom bar plot shows the total number of binding sites of the TFs. The right bar plot shows the number of regions that use the programs. The top and left dendrograms were computed by applying hierarchical clustering on the regulatory program matrix with Pearson correlation distance and average linkage. The program and TF labels are colored according to the hierarchical clusters. Selected groups of similar programs are labelled based on the major TFs participating in each program.

We found that the discovered regulatory programs are easy to interpret and reveal coherent functional groups of co-binding TFs. The programs discovered from K562 cells capture known sets of factors that interact with each other or function as a complex, such as the following:

- the master regulators GATA1, GATA2 and TAL1 (Cantor and Orkin 2002); and the enhancer-binding co-activator p300
- the transcriptional machinery Pol2, TBP, and TAF1; promoter-binding TFs such as E2F6 (Xu et al. 2007); and start site associated chromatin regulators such as PHF8 (Vermeulen et al. 2010), etc.
- CTCF, cohesin subunits RAD21 and SMC3, and ZNF143 (Bailey et al. 2015; Guo et al. 2012b)
- Pol3 transcriptional machinery (White 2011)
- AP-1 factors such as JUN/JUNB/JUND/FOS/FOSL (Chinenov and Kerppola 2001)
- MAF/BACH1/NFE2 (Kannan et al. 2012)
- MYC/MAX/USF/E2F6 (Blais and Dynlacht 2004)
- SPI1 ( also known as PU.1) and ELF1 (Bockamp et al. 1998)

To facilitate interpretation, we further cluster programs that are driven by similar sets of TFs into 23 program groups. The regulatory programs in the same group share the same set of main TFs, yet differ in some specific minor TFs. For example, in the CTCF group, programs 29 and 35 both contain the TFs CTCF, RAD21, SMC3, and ZNF143, Program 29 includes ARID3A and CEBPB, and Program 35 includes MAX, ZBTB7A, MYC, and YY1.

RPD is able to discover widely used regulatory programs as well as very specific programs. For example, the promoter, enhancer, and CTCF programs are each used in more than 10,000 regulatory regions. At the same time, small and specific programs such as ZNF274+TRIM28+SETDB1 (Program 48), which has been shown to bind at the 3’ ends of zinc finger genes and suppress their expression (Frietze et al. 2010), is used in only 63 regions.

The programs we discovered reveal context-dependent co-binding as reported in previous work. For example, we found that FOS mainly participates in five programs (Supplemental Fig. S3), which recapitulate four categories of FOS co-localization patterns reported previously (Xie et al. 2013). Although the fifth FOS category “AP1-HOT” does not directly correspond to a single program, it can be factored into AP1 and promoter programs that are mixed in those regulatory regions. In addition, we found that the category of FOS+NFYB can be further divided into FOS+NFYB+SP2 (Program 40) and FOS+NFYB+USF (Program 32).

We next compared factor co-occurrence in regulatory programs with published protein-protein interactions to see if the discovered programs were consistent with known factor interactions. In a published dataset of 33 protein-protein interactions among the 115 TFs in K562 cells (Xie et al. 2013), 22 of the TF physical interactions are captured as TF combinations described by the discovered regulatory programs (Supplemental Table S2).

Thus, RPD is sensitive enough to discover programs that are used in small number of regions and is able to recapitulate previous findings with a relatively small number of interpretable programs.

We then examined if the discovered regulatory programs were consistent with the chromatin state of the regulatory regions that use them. We annotated regulatory regions by DNase hyper-sensitivity, histone modifications, and the genome segmentation annotations derived from them (Hoffman et al. 2013; The ENCODE Project Consortium 2012). For the regulatory regions that utilize the same regulatory programs, we computed the fraction of the regulatory regions that overlap with these annotations (Fig. 3A). We found that for the regions using the same programs, the chromatin states of the regions are consistent with the functions of the TFs participating in the programs. For example, programs with master regulators of K562 cells and co-activator/co-repressors GATA1+GATA2+TAL1+p300+RCOR1+TEAD4 are used in regions that are annotated with enhancer chromatin state and are enriched with H3K4me1 histone modification, while the Pol2/promoter programs are used in the regions that are annotated with TSS chromatin state and are enriched with promoter-associated histone modifications such as H3K4me3, H3K9ac, and H3K27ac. Most regulatory programs are used by regulatory regions that are DNase hypersensitive, which is consistent with the preferential binding of TFs in open chromatin. Several programs used in the non-DNase hypersensitive regions are pre-dominantly repressive programs such as Program 48 (the combination of ZNF274, ZNF143, TRIM28, CBX3, SETDB1, and other factors) and Program 7 (the combination of ZNF143, TRIM28, CBX3 and SETDB1, but not ZNF274).

**Figure 3.**
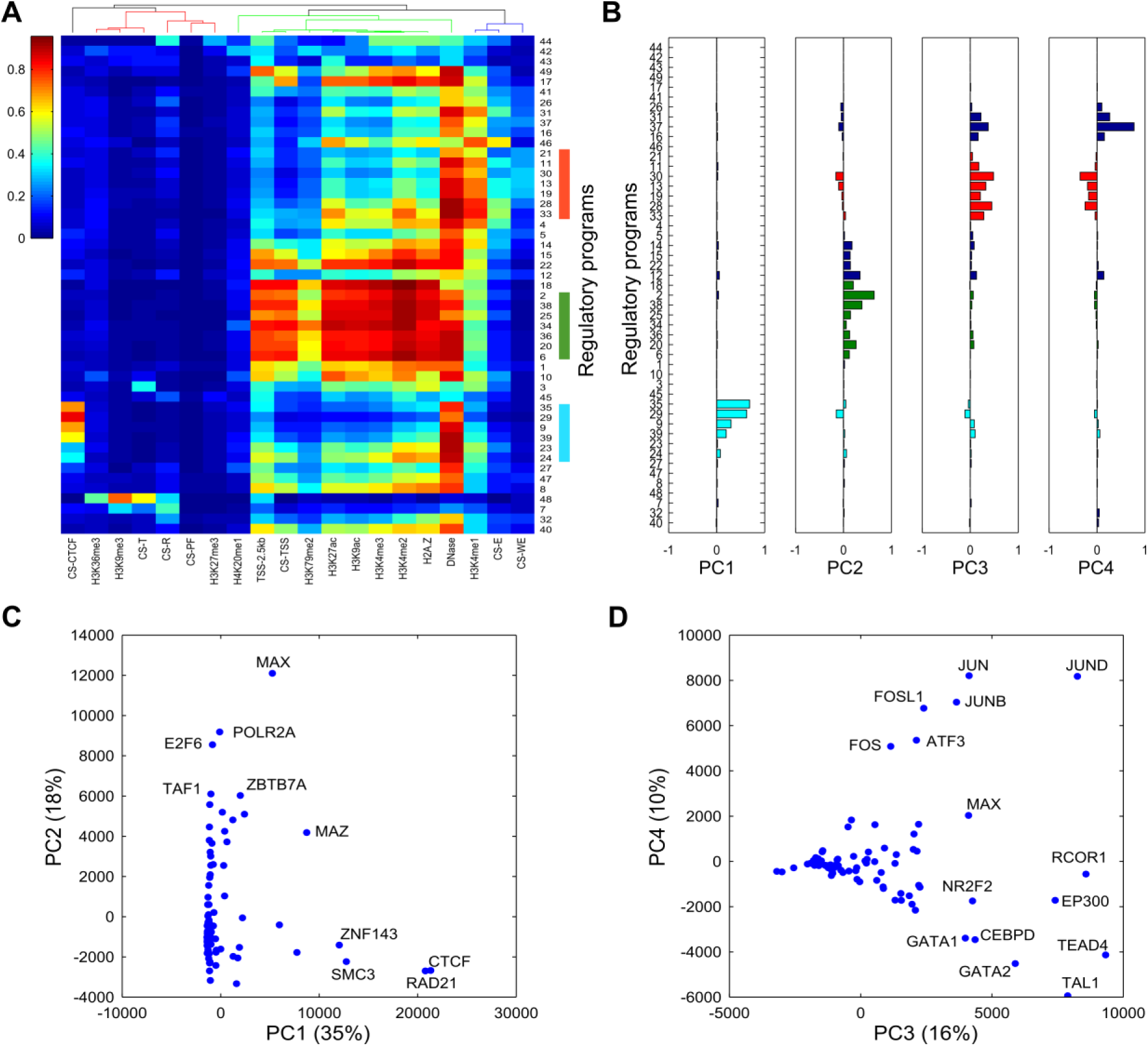
Epigenomic annotation and principal component analysis of the K562 regulatory programs. A) A heatmap showing the fraction of regions using the same regulatory programs (rows) overlapping the specific annotations (columns). The regulatory programs (rows) are ordered as in Figure 2. The top dendrogram was computed by applying hierarchical clustering on the fraction matrix with Pearson correlation distance and average linkage. B) Principal component analysis (PCA) was performed on the factor-program matrix. The PCA loadings of the regulatory programs for the first four PCs. C) The scatter plot shows the TF binding site counts projected on the principal components (PCs) 1 and 2. The percentage labels in the x and y axes represent the percentages of variance explained by the PCs 1 and 2, respectively. D) Similar to C). The TF binding site counts projected on the PC3 and PC4. CS: chromatin states; T: transcribed region; R: repressed region; PF: promoter flanking region; TSS: promoter region including TSS; E: enhancer region; WE: weak enhancer or open chromatin region.

We applied principal component analysis (PCA) to the program-TF matrix to reveal the structure and importance of the discovered regulatory programs. We found that programs are reduced to principal components (PCs) that correspond well with the major program groups (Fig. 3B-3D): the first PC is contributed mainly by the CTCF programs, explaining 35% of the total variance. The second PC is contributed mainly by the Pol2/promoter programs, explaining 18% of the total variance. The third and fourth PCs are contributed mainly by the p300/enhancer and AP-1 programs, explaining 16% and 10% of the total variance, respectively. In total, the first four PCs account for 79% of the total variance. These results indicate the dominant roles in genome-wide DNA binding of CTCF and cohesin, which have been suggested as key participants in shaping 3-dimensional genome structure (Phillips-Cremins et al. 2013), followed by promoter-binding factors and enhancer-binding factors. We found that the relative influence of the programs is not correlated with the number of factors in the programs because fewer factors contribute to the CTCF programs than to the promoter or enhancer programs.

### Many regulatory regions use more than one program

To understand the interplay among regulatory programs in regulatory regions, we next investigated the extent of multiple-program usage in regulatory regions. We found that 25,107 regulatory regions (~18%) use more than one program (Fig. 4A). For example, we found 3,742 regions that use both enhancer and AP-1 programs, 3,071 regions that use both promoter and CTCF programs, and 2,514 regions that use both promoter and enhancer programs. Interestingly, the regions that use both enhancer and promoter programs are annotated as either TSS/promoter or enhancer/weak enhancer chromatin states (Hoffman et al. 2013), with more than 63% of the regions being marked by both H3K4me1 (enhancer-related) and H3K4me3 (promoter-related) histone modifications (Supplemental Fig. S4). This represents a limitation of genome annotation methods that only assign a single label to a genome segment. Furthermore, a program associates with distinct types of regulatory regions when it co-occurs with different other programs. For example, the regions that use AP-1 programs co-occurring with enhancer programs are annotated as strong/weak enhancer chromatin state (Hoffman et al. 2013), AP-1 co-occurring with CTCF programs are annotated as CTCF state, and AP-1 co-occurring with promoter programs are annotated as promoter state (Fig. 4B).

**Figure 4.**
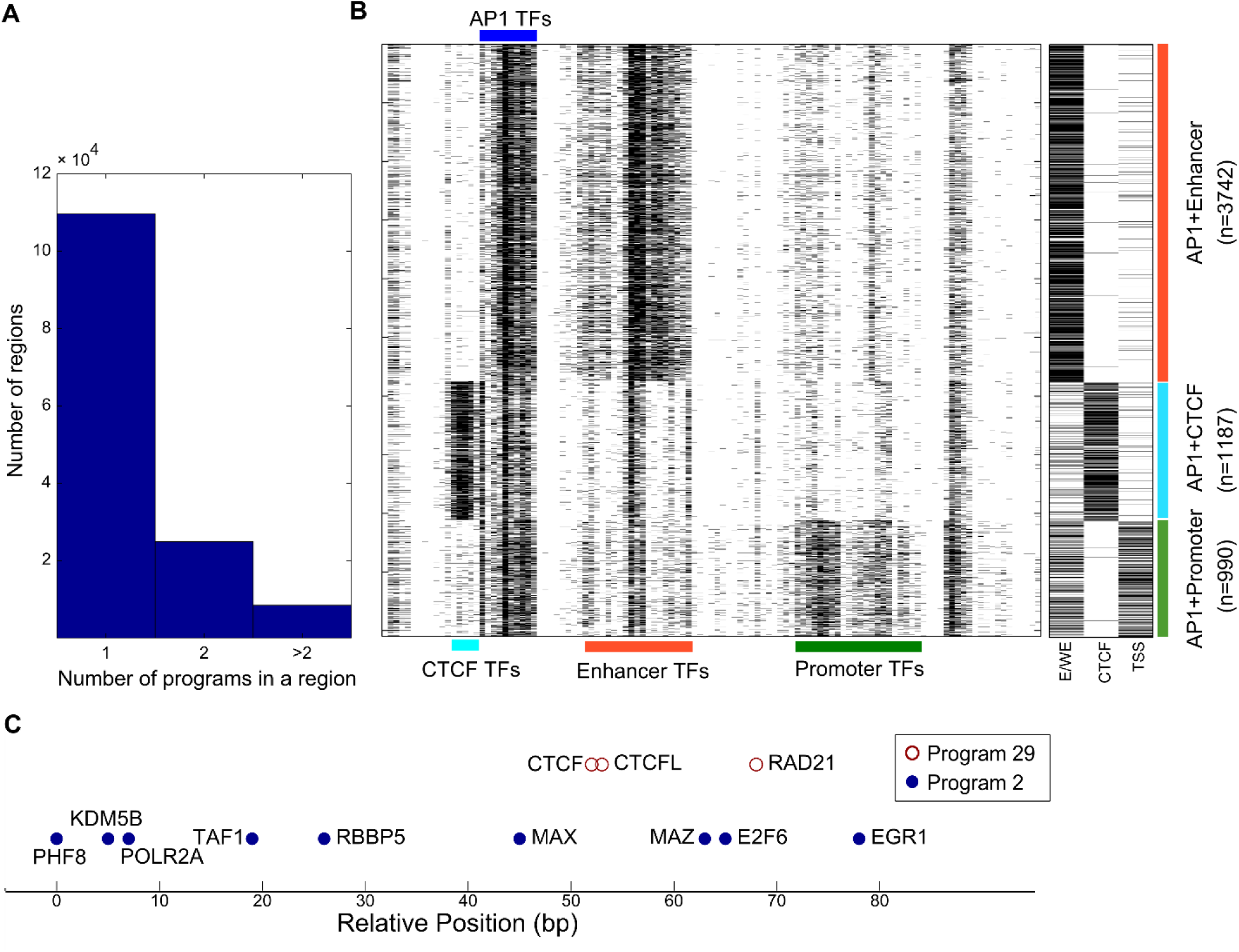
A large number of regulatory regions use multiple regulatory programs. A) A histogram of the number of regulatory programs used in a regulatory region. B) Co-occurring programs partition the regions use AP-1 programs into distinct functional categories. Left panel: A heatmap showing the region-TF binding matrix of the regions co-bound by AP-1 and enhancer programs, AP-1 and CTCF programs, and AP-1 and promoter programs. The TFs are in the same order as in Fig. 2. Right panel: A heatmap showing the chromatin state annotation of the same regions. E/WE: enhancer/weak enhancer state; CTCF: CTCF state; TSS: TSS/promoter state. C) An 80-bp region that contains binding sites assigned to a CTCF program (color in brown) and a promoter program (color in blue). The positions of the TF binding sites are called by GPS. Note that the positions of the sites assigned to different programs are mixed spatially.

To ensure that the discovery of multiple-program usage was not the result of inappropriate merging of proximal regulatory regions, we studied the positions of the binding sites that are assigned to different regulatory programs. In many cases, the binding sites assigned to distinct programs are spatially mixed. For example, in an 80bp region on chromosome 1, CTCF, CTCFL and RAD21 sites from a CTCF program are mixed with POL2, E2F6, MAX, EGR1, and other sites from a promoter program (Fig. 4C). Such co-occurrences between CTCF and promoter programs are consistent with previous findings that CTCF mediate long-range DNA-looping interactions between enhancers and promoter (Guo et al. 2012b), and that ZNF143 binds directly to the promoters and occupies anchors of chromatin interactions connecting promoters with distal enhancers (Bailey et al. 2015). Our analysis suggests that multiple-program usage is a prevalent aspect of regulatory activities in the cells and that it can be revealed by RPD.

### Regulatory program analysis identified distinct binding partners of NFY in different regions

In addition to identifying the global trends in the regulatory programs, we can use the combinatorial patterns for certain groups of TFs to generate hypotheses about TF interactions and then validate them with further analyses. For example, NFYA and NFYB (two components of NFY) both participate in Programs 32 and 40, together with the co-binding partner FOS (Fleming et al. 2013; Xie et al. 2013). Interestingly, NFY and FOS associate with USF1, USF2, ATF3, and MAX in Program 32, but with SP1 and SP2 in Program 40 (Fig. 5A). Because the mixed-membership nature of the topic model may allow these two programs to be both used in the same regulatory regions, we verified that these two programs are used predominantly in different regions. More specifically, we found 1227 regions that are bound by both NFY and USF2 but not by SP2, 1758 regions that are bound by both NFY and SP2 but not by USF2, and only 120 regions that are bound by NFY, SP2, and USF2 (Fig. 5B). The majority of the regions using the NFY+SP program are proximal regions, while most of the NFY+USF regions are distal regions. Furthermore, NFY and USF2 co-binding exhibits a strong spacing constraint, with 1227 co-bound regions exhibiting a 21-22bp spacing between NFY and USF2 motif supported sites. On the other hand, NFY and SP2 co-binding does not appear to have a specific spacing constraint. Thus, the observed co-occupancy and spacing constraints supports the binding of NFY and FOS with distinct combination of factors in different types of regulatory regions. Nf-y and Sp2 have been identified as co-binding partners that recruit each other to promoter regions in mouse embryonic fibroblasts (Völkel et al. 2015), and Nf-y and USF have been shown to co-bind in the promoter of mouse Miwi gene (Hou et al. 2012). Here we find that NFY+SP and NFY+USF co-binding occurs at thousands regions in human cells. Furthermore, our results show extensive co-binding of NFY+SP and NFY+USF in a mutually exclusive manner, suggesting the possibility of different DNA binding modes for NFY. These results highlight that systematically discovered regulatory programs may be used to generate specific hypothesis that can be tested with more detailed analysis.

**Figure 5.**
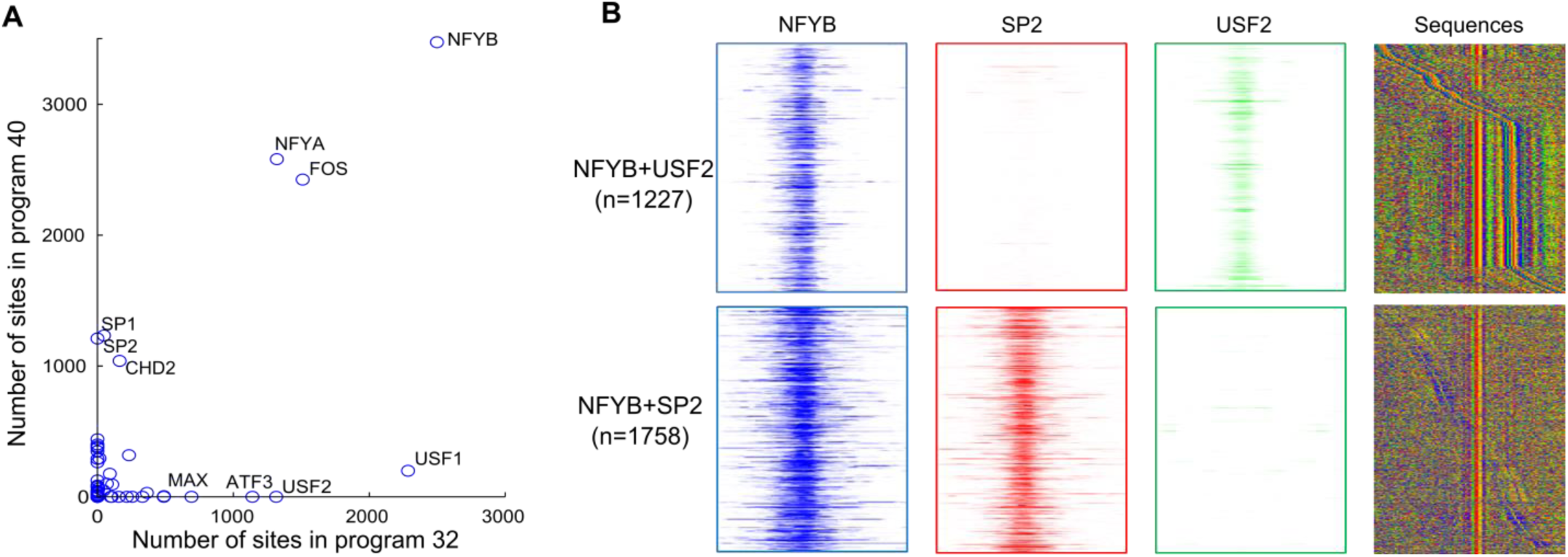
Regulatory program analysis identified distinct binding partners of NFY in different regions. A) NFY (NFYA and NFYB) and FOS participate in two distinct programs. They co-bind with SP1 and SP2 in program 40 while in program 32, they co-bind with USF1 and USF2, ATF3 and other factors. The scatter plot shows the number of binding sites of each factor in the two programs. B) NFYB binds exclusively with USF2 or SP2, each with different spacing constraints. The first 3 columns show ChIP-seq read enrichment of NFYB, SP2, and USF2 in a 1-kbp window around the NFYB binding sites. The 4th column shows sequence plots in a 100bp window around the NFYB binding sites. The regions are centered at the NFYB binding sites and sorted by the distance between NFYB and USF2 or NFYB and SP2, respectively. The sequence motifs for USF2 and SP2 can be observed around the NFYB motifs. Green, blue, yellow, and red indicate base A, C, G, and T.

### The combinatorial rules of direct/indirect binding

We next compared the binding partners of TFs when they bind directly (motif present) and indirectly (motif absent) to the genome. Here we define direct binding as the binding of a TF at sites that contain the cognate motif of the TF, and indirect binding as binding at sites that do not contain the cognate motif. Previous studies have found that many ChIP-seq binding sites of sequence-specific TFs do not contain the cognate motif of the TFs, suggesting that binding may be indirect through the interaction with co-binding TFs (Farnham 2009). Understanding the combinatorial patterns of direct binding versus indirect binding may reveal the co-binding TFs that facilitate indirect binding.

For each sequence-specific factor X, the binding sites were divided into two groups, dX sites (direct binding) where X motif is present and iX sites (indirect binding) where X motif is not present. These two groups were then treated as two separate factors. For the K562 dataset, this motif-based division expanded the total number of the factors to 167, with 52 pairs of direct and indirect binding “factors”. Applying RPD to these data, we discovered 54 regulatory programs with the expanded set of direct and indirect factors (Supplemental Fig. S4, Supplemental Table S1) to investigate the presence of co-binding factors that are specific to direct or indirect binding. For example, previous work reported that FOS co-localizes with NFYB (Xie et al. 2013). Our analysis showed that this co-localization mostly occurs between indirect FOS binding and both direct and indirect NFYB binding (Programs 20 and 24) (Fig. 6A). To more systematically investigate the indirect binding of factors, we compute the pairwise correlation between all the 52 direct binding factors and all the 52 indirect binding factors (Fig. 6B). A high correlation would indicate that the indirect binding factor likely associates with the corresponding direct binding factor. We found several groups of such associations: indirect binding of FOS with direct binding of NFY+SP1+SP2, indirect binding of E2F6 with direct binding of MYC+MAX+BHLHE40+USF+MXI1+YY1, and indirect binding of ATF3 with FOS+FOSL1+JUNB+JUND+JUN.

**Figure 6.**
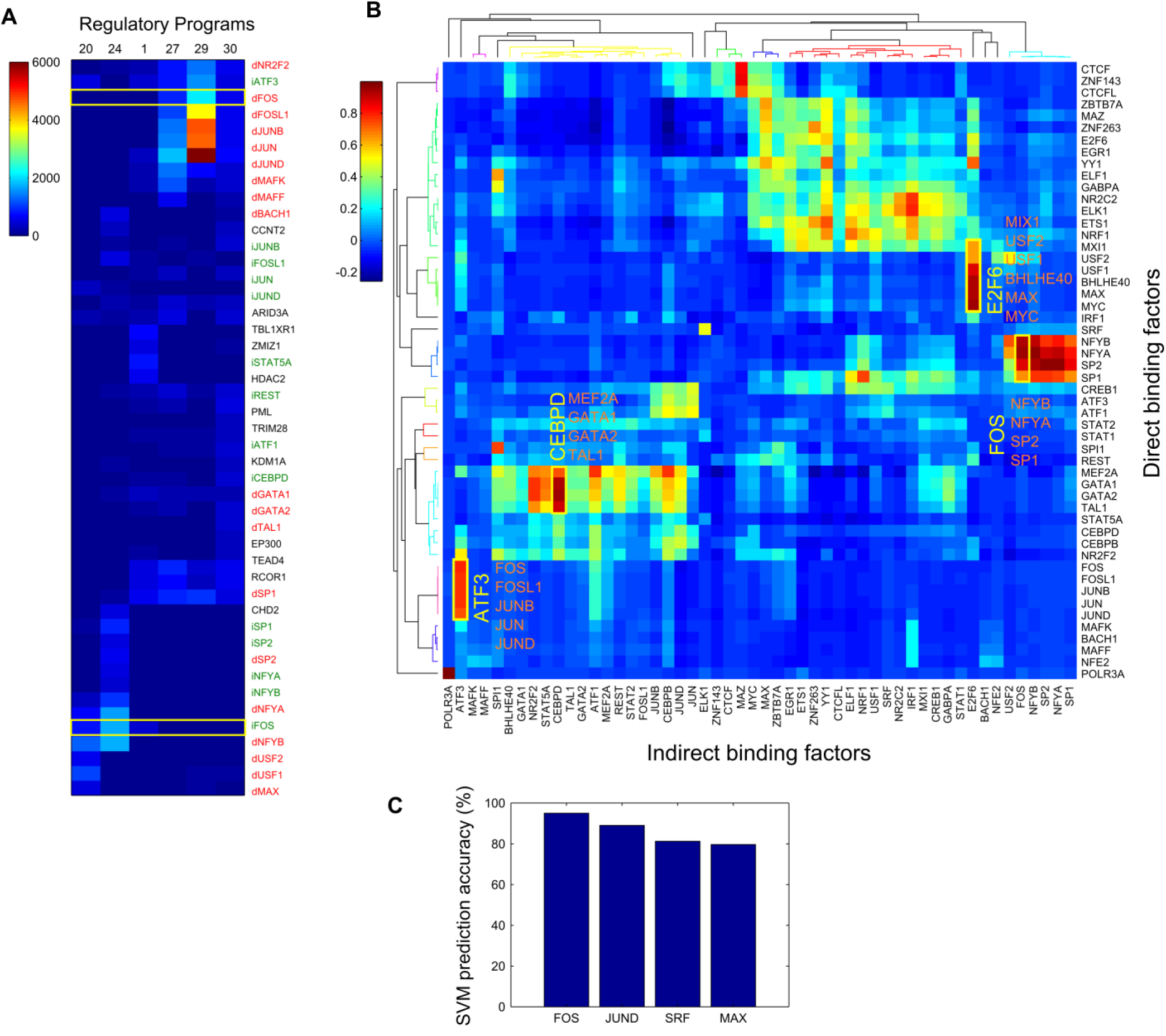
Direct/indirect binding of some TFs can be explained by the specific combination of co-binding TFs. A) FOS co-binds with different set of TF partners when binding directly or indirectly. A subset of the regulatory program matrix that involves FOS binding are shown. Each cell in the heatmap represents the TF binding site count of a TF (row) in a regulatory program (column). dTF (in red text) represents the direct binding sites (motif present) of the TF. iTF (in green text) represents the indirect binding sites (motif absent) of the TF. B) RPD discovered specific combinations of TF indirect and direct binding. Sequence-specific TF binding sites are divided into direct binding sites and indirect binding sites based on whether a site contains a motif of the TF. RPD is applied to the expanded set of TFs to discover regulatory programs. The heatmap shows the correlation between direct binding and indirect binding of 48 TFs based on their program participation. The top and left dendrograms were computed by applying hierarchical clustering on the correlation matrix with Pearson correlation distance and average linkage. Yellow boxes label some combinations of TF indirect binding (yellow text) and direct binding (orange text). C) Direct/indirect binding of some TFs can be predicted with high accuracy using the specific combination TFs binding in the same regions. A support vector machine (SVM) classifier was trained to predict whether a TF binds directly or indirectly using the binding of the other TFs. The SVM was trained on chromosome 1 and tested on the rest of the genome.

Furthermore, we observed that the direct binding sites and indirect binding sites of some factors participate in very different programs. For example, direct FOS binding co-occurs with AP-1 factors (Program 29) and MAF+BACH1+NFE2 (Program 27), while indirect FOS binding co-occurs with NFY+SP1+SP2 (Program 24) or NFY+USF1+USF2 (Program 20) (Fig. 6A).

With the observation that direct and indirect binding sites of some factors associate with different combination of factors, we reasoned that it would then be possible to predict whether a sequence-specific TF binds DNA directly or indirectly based on the proximal binding of other TFs. To test this hypothesis, we trained a support vector machine (SVM) classifier to predict whether a TF binding site is a direct or indirect site using the proximal binding of other TFs in the region. For each sequence-specific TF, we trained an SVM with data on chromosome 1, and then tested on the other chromosomes. For factors such as FOS, JUND, SRF, and MAX, the SVMs predict the direct/indirect binding of the factors with 80-95% accuracy (Fig. 6C). Consistent with the SVM prediction performance, the co-binding factor combinations are different when these factors bind directly or indirectly (Supplemental Fig. S6).

### Cell-type-specific programs

We next investigated if we could observe cell-type specific regulatory programs and cell-type common regulatory programs. We used ChIP-seq data of 56 TFs that were profiled in both K562 and GM12878 cells from the ENCODE Project (The ENCODE Project Consortium 2012). Following a previous approach (Xie et al. 2013), we constructed the co-binding regions in each cell type separately (~105,000 regions in K562 and ~91,000 regions in GM12878) and then combined the data from all regions from both cell types for RPD analysis. RPD discovered 48 programs that describe the binding of the 56 factors in K562 and GM12878 cells (Fig. 7A, Supplemental Table S1). To aid interpretation of the programs we clustered them into 16 program groups. Interestingly, the promoter-associated programs and CTCF-associated programs are each clustered into one program group, while the enhancer-associated programs are clustered into two groups.

**Figure 7.**
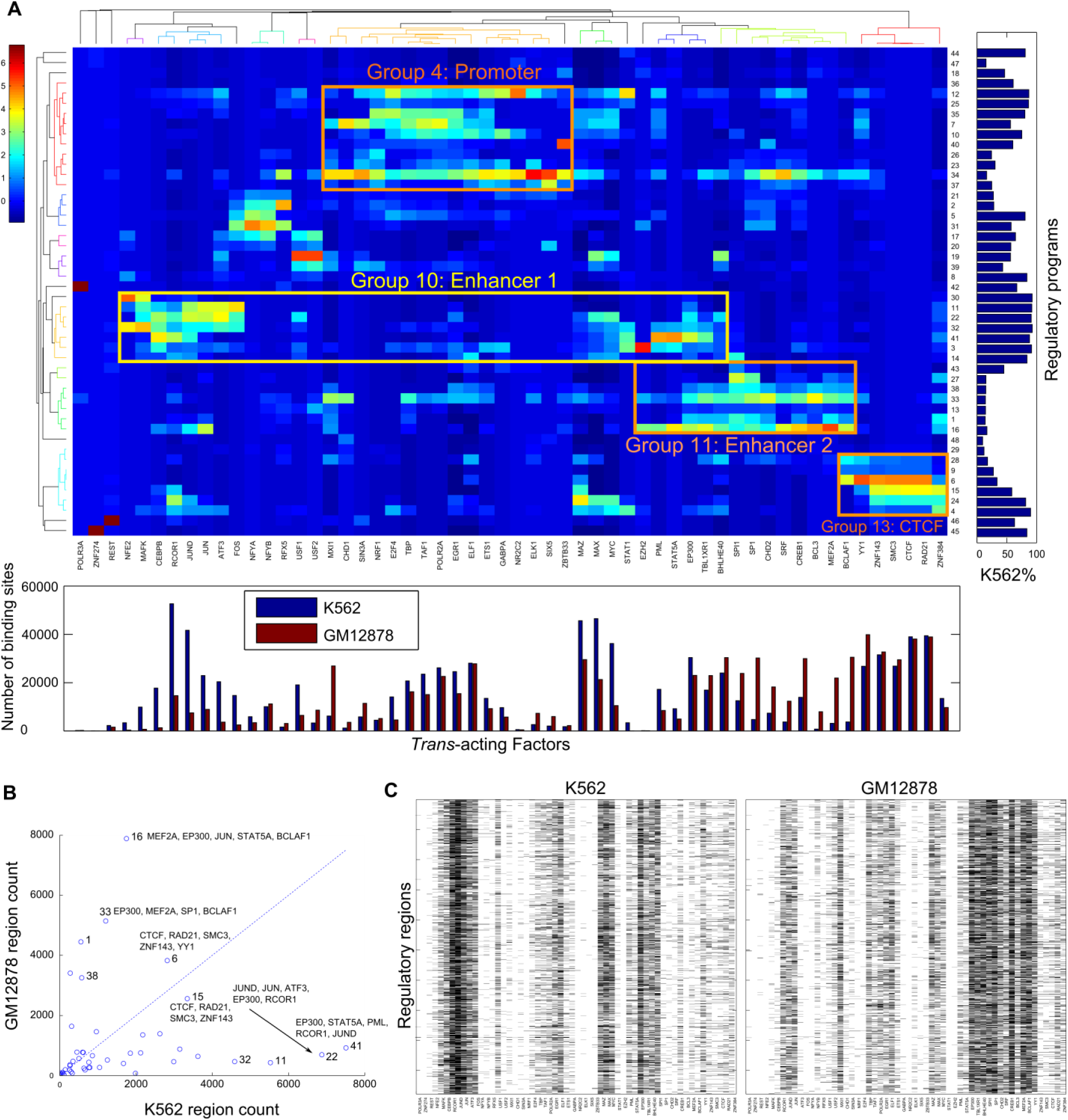
Common and cell-type-specific regulatory programs. A) RPD was applied to ChIP-seq binding sites of 56 TFs in human K562 cells and GM12878 cells. Each cell in the heatmap represents the z-score of TF binding site count (standardized along the columns) of a TF (column) in a regulatory program (row). The bottom bar plot shows the total number of binding sites of the TFs. The right bar shows the percentage of the sites contributed from K562 data for the programs. The top and left dendrograms were computed by applying hierarchical clustering on the regulatory program matrix with Pearson correlation distance and average linkage. Groups of similar programs are labelled based on the major TFs participating each program. B) A scatter plot showing that the cell-type-specific programs are preferentially used in the differentially bound regions in K562 cells and GM12878 cells. C) Distinct cell-type-specific programs are used in the same genomic regions in different cell types. A set of 1956 regions that are bound by K562 enhancer program factors in K562 (left panel) and by GM12878 enhancer program factors in GM12878 cells (right panel). The TFs are in the same order as in A).

To investigate the degree of cell type specificity of the discovered regulatory programs, we computed the fraction of the binding regions that are contributed from the K562 data or from the GM12878 data for each program. We found that some combinations of factors are mainly used in K562 cells and others are mainly used in GM12878 cells even though all of the 56 factors bind regulatory regions in both cell types (Fig. 7A). In particular, the enhancer program groups are preferentially used in only one of the cell types. In K562 cells p300 co-binds with JUND, JUN, RCOR1, ATF3, FOS, CEBPB, MAFK, MAX, and MYC (Program group 10), but in GM12878 cells p300 co-binds with MEF2A, SP1, SPI1, BCLAF1, BCL3, and BHLHE40 (Program group 11). Both cell types share other program groups, such as the promoter and CTCF program groups. In the regulatory regions that are bound in K562 or GM12878 cells but not bound in both cell types, cell-type-specific programs are used preferentially by one cell type, while shared programs are used in both cell types (Fig. 7B).

We next investigated if regulatory regions that are bound in both cell types use the same or different regulatory programs. Out of 50,910 regions that are bound in both cell types, we found that 1,956 regions use a different enhancer program in the two cell types (Fig. 7C). Program group 10 is used in the K562 cells and program group 11 is used in the GM12878 cells. Thus although these regions are bound in both cell types and may act as enhancers, as suggested by the binding of transcriptional co-activator p300, they are bound by cell-type specific combinations of factors in K562 and GM12878. In addition, we found 1,312 regions that are bound by CTCF program factors in both K562 and GM12878 cells, and also bound by the K562 enhancer program factors in K562 cells, suggesting the usage of K562-specific enhancers in these CTCF/cohesin bound regions. In summary, important differences can exist between the set of regulatory programs that bind the same regulatory regions in distinct cell types.

## Discussion

We have demonstrated that RPD’s systematic dissection of combinatorial binding of *trans*-acting factors (TFs) can provide important insights into the mechanisms of gene regulation not available with previous hard-clustering-based methods. Our approach models the modular usage of regulatory programs in a regulatory region, and summarizes high-dimensional binding data into a relatively small number of compact and coherent programs. The programs discovered are easy to interpret individually and as a whole, uncovering specific partners of individual TFs as well as the global structure of the combinatorial binding patterns.

We have found that the regulatory programs discovered by RPD capture key aspects of global combinatorial binding patterns and provide a resource for generating new hypotheses for TF interactions. Our analysis reveals extensive TF combinatorial binding and multiple-program usage in tens of thousands of regulatory regions. We discovered that the direct and indirect DNA binding of certain TFs is associated with distinct sets of binding partners and that such information can predict whether the TF binds directly or indirectly with high accuracy. Finally, our analysis discovered cell-type-specific programs and shared programs, and that thousands of regulatory regions use different programs in different cell types.

Our approach is able to identify regulatory regions that are bound by multiple regulatory programs. Understanding multiple-program usage is important because specific gene regulation in the cells is initiated by the interactions among a complex group of TFs that are organized as distinct programs. Previous methods have modeled TF binding at a given regulatory region with a single regulatory program and thus are not able to capture the complexity of modular combinatorial binding. Our analysis reveals the widespread usage of multiple modular regulatory programs in tens of thousands of regions. With a larger number of additional TFs being assayed by large-scale efforts such as the ENCODE project (The ENCODE Project Consortium 2012), we expect that RPD will be useful in revealing the complexity of combinatorial binding in these data.

One potential limitation on studying combinatorial TF binding from ChIP-seq data is the relative scarcity of high quality antibodies. To expand RPD combinatorial binding analysis to more TFs or to cell types that do not have sufficient ChIP-seq data, one strategy is to augment or replace ChIP-seq data with TF binding predicted from DNase-seq (Thurman et al. 2012) or ATAC-seq (Buenrostro et al. 2013) data and TF motif information (Pique-Regi et al. 2011; Sherwood et al. 2014).

## Methods

### Data and preprocessing

ChIP-seq data for the TFs and corresponding controls were downloaded from the ENCODE project website http://hgdownload.cse.ucsc.edu/goldenPath/hg19/encodeDCC/. Fastq files were aligned to hg19 genome with Bowtie (Langmead et al. 2009) version 0.12.7 with options “-q --best --strata -m 1 -p 4 --chunkmbs 1024”. GEM (Guo et al. 2012a) was used to call binding events with default parameters using the aligned reads of TF ChIP-seq experiments and the corresponding control experiments. GEM produces two set of binding site calls for each dataset: GPS binding calls without motif information and GEM binding calls with motif information. The binding calls overlapping with the ENCODE blacklist regions (http://hgdownload.cse.ucsc.edu/goldenPath/hg19/encodeDCC/wgEncodeMapability/wgEncodeDacMapabilityConsensusExcludable.bed.gz) were excluded for this analysis.

### Construct co-binding regions

GPS binding calls of all the factors in a given cell type were pooled together to construct the co-binding regions. Each binding call was expanded +/−50bp from the summit position. Then overlapping binding calls were merged to form non-overlapping co-binding regions. For this study, only co-binding regions with three or more binding calls were used for subsequent analysis. The expansion length of 50bp was used to construct co-binding regions because the spatial resolution of TF ChIP-seq binding calls is about 30-50bp. Using 100bp expansion length to construct co-binding regions gives similar results. From the K562 co-binding regions (n=142,962), we construct a Region-TF matrix (142,962x115) that contains the number of binding sites of each TF in each region.

### Topic model

The hierarchical Dirichlet processes (HDP) topic model was used in this study because it automatically determines the number of the topics from the data. A C++ implementation of HDP was downloaded from http://www.cs.columbia.edu/~blei/topicmodeling_software.html. The parameters used were “--eta 0.1 --max_iter 2000”. Eta is the hyperparameter for the topic Dirichlet distribution. We tested different eta values (0.01, 0.05, 0.1, 0.5 and 1) and the results were similar. We chose eta to be 0.1 to encode our assumption that each topic contains only a few TFs. The HDP inference procedure typically converged at about 1000 iteration. We ran the HDP with 3 different random seeds for 2000 iterations and used the run that had the highest data likelihood reported by the HDP. The input to the HDP are the TF binding site counts in the co-binding regions. Each region is treated as a document and the TF sites as words in the documents. The output of the HDP includes the program-TF matrix and the program assignment of each TF binding site.

For the program-TF matrix, each column vector (TF participation vector) describes the distribution of the TF binding sites across all the programs, and each row vector (program vector) represents the number of binding sites contributed by each TF to the program. We compute a z-score for each TF vector. A TF is considered to participate a program if the z-score of the TF-program pair is larger than 1. Similarly, we compute a z-score for each program vector. A TF is considered to be a “main driver” of the program if the z-score of the TF-program pair is larger than 1. Each program is labeled with the names of the main TF drivers, which are ranked by their z-scores. To facilitate interpretation of the programs, the program-TF matrix was clustered into program groups using hierarchical clustering with Pearson correlation distance and average linkage. The cutoff distance for clustering is 0.5.

The region-program assignment table assigns each TF binding site from each region to one of the programs. The assignment table was summarized into a region-program matrix where each element of the matrix represents the number of the TF binding sites in a region that are assigned to a particular program. A program is considered as being used in a particular regulatory region if 1) at least three binding sites in the region are assigned to the program and 2) the z-score of the site count for the program in the region is larger than 1.

### Compare HDP with k-means clustering and NMF

To test the ability of RPD, k-means clustering, and NMF to accurately capture TF-TF correlations that are present in the binding data, we applied all three approaches to the same set of K562 cell ChIP-seq binding data to discover the same number of programs (k=49). We applied k-means clustering and NMF on the K562 Region-TF binding matrix using the MATLAB software (MATLAB and Statistics Toolbox Release 2012a, The MathWorks, Inc., Natick, Massachusetts, United States). For k-means clustering, Euclidean distance is used as the distance metric. The cluster number (i.e. rank for NMF) was set to be k=49 and k=100 to compare with the HDP topic model with 49 topics. We refer the k-means clusters, NMF components and HDP topics as the programs. To compare the three approaches, we first computed the pairwise TF correlation scores using the original program-TF matrix, or the program-TF matrices derived from these three methods. Then we computed the correlation between the pairwise TF correlation scores from the original region-TF matrix and those from the three derived program-TF matrices. Results are presented in Fig 1B and 1C. We also compared topic model programs and k-means clusters by computing their Pearson correlation for the program vectors vs. the cluster vectors. Results are presented in Fig S1.

### Protein-protein interaction and epigenomic annotation of regulatory programs

The protein-protein interaction derived from IP-MS Data for K562 cells (Xie et al. 2013) was downloaded from http://www.cell.com/cms/attachment/2021777707/2041662737/mmc1.xls. For the 33 direct physical interaction pairs that contain the TFs in our study, we considered an interaction as rediscovered by RDP if the two TFs are both the “main drivers” in a same regulatory program.

DNase hyper-sensitive open chromatin peak calls, histone modification peak calls, and the combined genome segmentation annotations were downloaded from http://ftp.ebi.ac.uk/pub/databases/ensembl/encode/integration_data_jan2011/. The co-binding regions were annotated with the epigenomic annotations if they overlap at least 1bp. For each regulatory program, we identified the regulatory regions that use the program and computed the fractions of these regions that overlap with the annotations.

### Direct vs. indirect binding analysis

For the direct vs. indirect binding analysis, GEM binding calls were used for sequence-specific binding factors that the GEM motifs can be verified. The positional frequency matrix of top ranked motif reported by GEM was compared against known motifs of the same factor in the public databases using STAMP (Mahony et al. 2007), as previously described (Guo et al. 2012a). For the factors that a database match for the top motif is found, the GEM binding calls were divided into direct and indirect binding sites based on whether the binding sites contain a motif match of the TF. The direct and indirect binding sites were treated as separate factors for topic modeling analysis. For the non-sequence-specific factors and sequence-specific factors that the top GEM motif does not match the known database motifs of the factor, GPS binding calls were used. All the GEM and GPS binding calls were then pooled together to construct the co-binding regions. In total, 159,204 co-binding regions with binding sites from 167 “factors” were constructed. Applying RPD, we discovered 54 programs. The correlation between direct and indirect binding factors were computed using the TF vectors in the program-TF matrix.

### Predict direct/indirect binding using SVM

We trained a support vector machine (SVM) classifier to predict whether a TF binding site is a direct or indirect site using the proximal binding of other TFs in the co-binding region. We used the SVM software in the MATLAB software (MATLAB and Bioinformatics Toolbox Release 2012a, The MathWorks, Inc., Natick, Massachusetts, United States). For each sequence-specific TF, we trained the SVM with data on chromosome 1 and then tested on the other chromosomes. More specifically, using the region-TF matrix (159,204 x 167), we took the rows that contained either direct or indirect binding sites of the TF, used the columns corresponding to the direct or indirect binding of the TF as the prediction and the rest of the columns (binding of the other TFs) as the features. We tested linear kernel and Gaussian Radial Basis Function (RBF) kernel and found that the accuracy is higher when using the RBF kernel. The parameters rbf_sigma and boxconstraint for the svmtrain() function were determined by grid search using 5-fold cross-validation.

### Cross-cell-type analysis

For the cross-cell-type analysis, if there were multiple datasets for the same factor, we chose the datasets that were produced by the same lab, using the same antibodies, or had similar number of binding calls. The co-binding regions were constructed separately for K562 and GM12878. The data from both cell types were then concatenated for topic model analysis. Thus we can learn TF co-binding relationships that are shared across cell types but still keep track of the cell type origin of the TF sites and the regions.

## Acknowledgements

We thank Tatsunori Hashimoto and Haoyang Zeng for insightful comments and assistance in analysis. This work was supported by National Institutes of Health (grant 1U01HG007037-01 to D.K.G).Author contributions: Y.G. conceived the project, developed the method, analyzed the results, and wrote the manuscript. D.K.G. supervised the project and edited the manuscript. Both authors read and approved the final manuscript.

